# Variation in susceptibility determines the strength of observed community-level dilution effects

**DOI:** 10.64898/2026.11.24.769010

**Authors:** Megan C A Tanlimco, Daniel Pelletier, Leithen K M’Gonigle, Arne O Mooers, Ailene MacPherson

## Abstract

Understanding how pest abundance responds to biodiversity is critical in human-managed landscapes. Biodiversity is said to exhibit a dilution effect when the relationship is negative, but the general conditions under which this occurs (versus its converse, an amplification effect) are still contested. Using data on 52 pest species across 26,549 North American forest plots we document how the dilution effect is impacted by a pest’s host breadth. Further, we show the importance of considering how pest prevalence is measured. When pest prevalence is calculated across all trees in a forest plot, the dilution effect is stronger for specialist pests than for generalist pests, as expected. However, when prevalence is calculated only for trees known to be hosts of a pest, the dilution effect is stronger for generalist pests. A multi-host epidemiological model illustrates that simply ignoring locales without evidence of disease can give rise to these contrasting patterns. Given their impacts, it will be important to consider the effects of both a pest’s host breadth and methodology when predicting and assessing the impact of changing biodiversity on pest prevalence.

## 1 Introduction

A negative relationship between pest or disease prevalence and biodiversity, the dilution effect (Keesing et al., 2006), is a potentially important ecosystem service (Young et al., 2017). However, it is also contested: its converse, amplification, is also possible, and both are specific outcomes of the general disease diversity relationship, or DDR (Keesing et al., 2006). Even in well-studied and economically important ecosystems such as working forests, increased tree diversity has been linked to both decreased and increased pest prevalence (see citations in Gougherty and Davies, 2024). Such variation in the effects of biodiversity on pests can be due to ecological context (Halliday et al., 2020), pest host breadth (see, e.g., Jactel et al., 2021), and, importantly, how prevalence is measured.

Most generally, the dilution effect is possible (but not guaranteed) if two ecological conditions are met (Keesing and Ostfeld, 2021). First, potential host species must differ in how susceptible they are to the pest or pathogen (from here on in, we refer to pests) and/or how well they transmit pests to other host individuals. In other words, they must vary in competence. Second, increased biodiversity must decrease transmission from more competent hosts. This latter condition can occur, inter alia, via ”host regulation,” when the density or frequency of competent hosts decreases at high levels of biodiversity due to factors such as interspecific competition, which leads to reduced transmission. Under the assumption that low biodiversity communities are dominated by higher competence hosts and that high biodiversity communities favour lower competence hosts, we expect to see dilution effects by host regulation associated with a decrease in community-averaged competence.

Empirical studies do tend to find a negative relationship between community-averaged competence and biodiversity, especially in cases of biodiversity loss (Halliday et al., 2020; Johnson et al., 2013; Rosenthal et al., 2021). One explanation for this relationship is that fast life-history species are more competent hosts because they invest less in defenses, but are also less susceptible to local extirpation and, thus, are not the species that are lost when diversity drops (Joseph et al., 2013). Both experimental (Liu et al., 2018) and simulation (Ostfeld and LoGiudice, 2003) studies have shown that communities that differ in their diversity due to random species loss have weaker (or absent) dilution effects compared to communities that differ based on realistic or observed patterns of species loss, and this relationship between biased species loss and the dilution effect is supported by comparative data (Halliday et al., 2020).

Pest biology may also matter. Theory suggests that specialist pest species should exhibit a stronger dilution effect than generalists (Collins et al., 2020; Keesing and Ostfeld, 2021). This prediction is an extension of the necessary condition that dilution requires that competence varies among potential hosts of a pest (Keesing and Ostfeld, 2021). Suppose that we begin in a low biodiversity community dominated by hosts with high competence. If increasing biodiversity introduces host species with lower competence, the extent to which pest transmission and prevalence decreases depends on the degree to which host competence varies. On one extreme, a specialist pest that strictly uses one host species will experience great variation in competence among potential host species: their one host species will be highly competent while other species would have zero competence. This would produce the highest possible decrease in transmission and prevalence overall as biodiversity increases. On the other extreme, a generalist pest where all potential hosts are equally competent would not experience any effect as biodiversity increases. However, connecting this prediction to empirical tests requires an understanding of the assumptions and effects associated with various methodological decisions and limitations in studying the dilution effect.

The dilution effect can be assessed at the host species level, generating pest prevalence estimates for each host species individually, or at the community level, aggregating prevalence (fraction of affected trees) across multiple host species (Rosenthal et al., 2021). While these (and other) approaches can all lead to dilution effects, different mechanistic assumptions and interpretations apply to them (Rosenthal et al., 2021; Roberts and Heesterbeek, 2018). For instance, we would expect community-level prevalence to drop as biodiversity increases due to how extirpation risk and defenses correlate with life-history, but have less clear expectations if prevalence is measured for individual host species. Furthermore, prevalence at the community level can be quantified using different criteria for which species are included in the denominator. Prevalence could be calculated only across species known to be suitable hosts for the focal pest, which would quantify the extent to which the pest is filling its niche (e.g., Rosenthal et al., 2021). Alternatively, prevalence could be considered across a broader community using a denominator that includes the total number of both host and non-host species, such as all the tree species present in a forest (e.g., Gougherty and Davies, 2024). This approach quantifies how much a pest is affecting the chosen community as a whole.

In recent decades, accounting for user detection bias has become nearly mandatory in statistical analyses of species occurrence and abundance (e.g., occupancy models and their derivatives are now commonplace; MacKenzie et al., 2006). A similar problem likely exists here: individual pathogens are more likely to be studied in places where they are known to occur, such that real species’ absences are likely biased low, if they are recorded at all. For example, the forest plot data we consider below are missing pest species’ absences altogether. Such a bias in the underlying data will almost certainly affect any estimated DDR: for instance, the DDR should depend heavily on the frequency and composition of plots where any given pest is *not* detected.

In their study on the role of host evolutionary history in the dilution effect, Gougherty and Davies (2024) predict, as suggested by theory, that specialist forest pests would be subject to stronger dilution effects than generalists. Using the estimated DDR for a select subset of specialist and generalist species, they report that specialists and generalists both can be subject to strong or weak dilution effects and suggest no apparent relationship between generalism and the disease diversity relationship; however they do not formally test the prediction. Importantly, their analysis calculated community-level prevalence across *all* tree species; considering individual host species or only known host species could change their conclusions.

Here, we use the same forest pest dataset as Gougherty and Davies (2024) to explicitly test the hypothesis that specialists are more sensitive to the dilution effect than generalists. In our analyses and interpretation, we focus on two of the factors highlighted above: first, we compare patterns using two community-level interpretations of the dilution effect, one which assesses prevalence among only a pest species’ known tree host species and one that follows Gougherty and Davies (2024) by considering prevalence across all sampled tree species. We also consider the potential effects of bias in recorded species’ absences. To understand the possible mechanisms underlying our result, we develop and analyze a simple multi-host epidemiological model. This model suggests that the effects of (1) variance in a pest species’ host susceptibilities and (2) site selection bias can combine to explain variable effects of biodiversity on pest prevalence, both in our dataset and more generally. Given the potential importance of the dilution effect in mitigating pests and disease, resolving or accounting for these variable findings in future work is critical.

## 2 Results

### 2.1 Empirical Results

To estimate the disease-diversity relationship for individual pest species, we fit a Beta-Binomial model with species-specific effects of Shannon diversity to plot-level pest prevalence data. We included plot Shannon diversity and total number of either all trees or host trees as predictors of the number of affected trees for a focal pest in each plot. We refer to this model as the “Species-specific slopes model.” For a Beta-Binomial model, one must specify the number of “successes” and the number of “trials” for each data point. Here, the number of successes is the number of trees affected by a particular pest in a plot. For the number of trials, we conduct analyses considering each of two community definitions: 1) the number of trees in the plot that are a known host species of that pest species or 2) the total number of trees in the plot, regardless of whether or not they are known hosts for that pest species. These two definitions correspond to calculating prevalence across 1) known host trees (number of affected trees/number of host trees) or 2) all trees in the forest community (number of affected trees/total number of trees). We found that the choice of prevalence measure qualitatively determines the estimated disease-diversity relationship. Specifically, when we assessed prevalence across known host trees, species showed a mix of positive and negative disease-diversity relationships (Fig. 1A), whereas when we assessed prevalence across all trees, all but one pest species showed a negative disease-diversity relationship (Fig. 1B).

**Figure 1:**
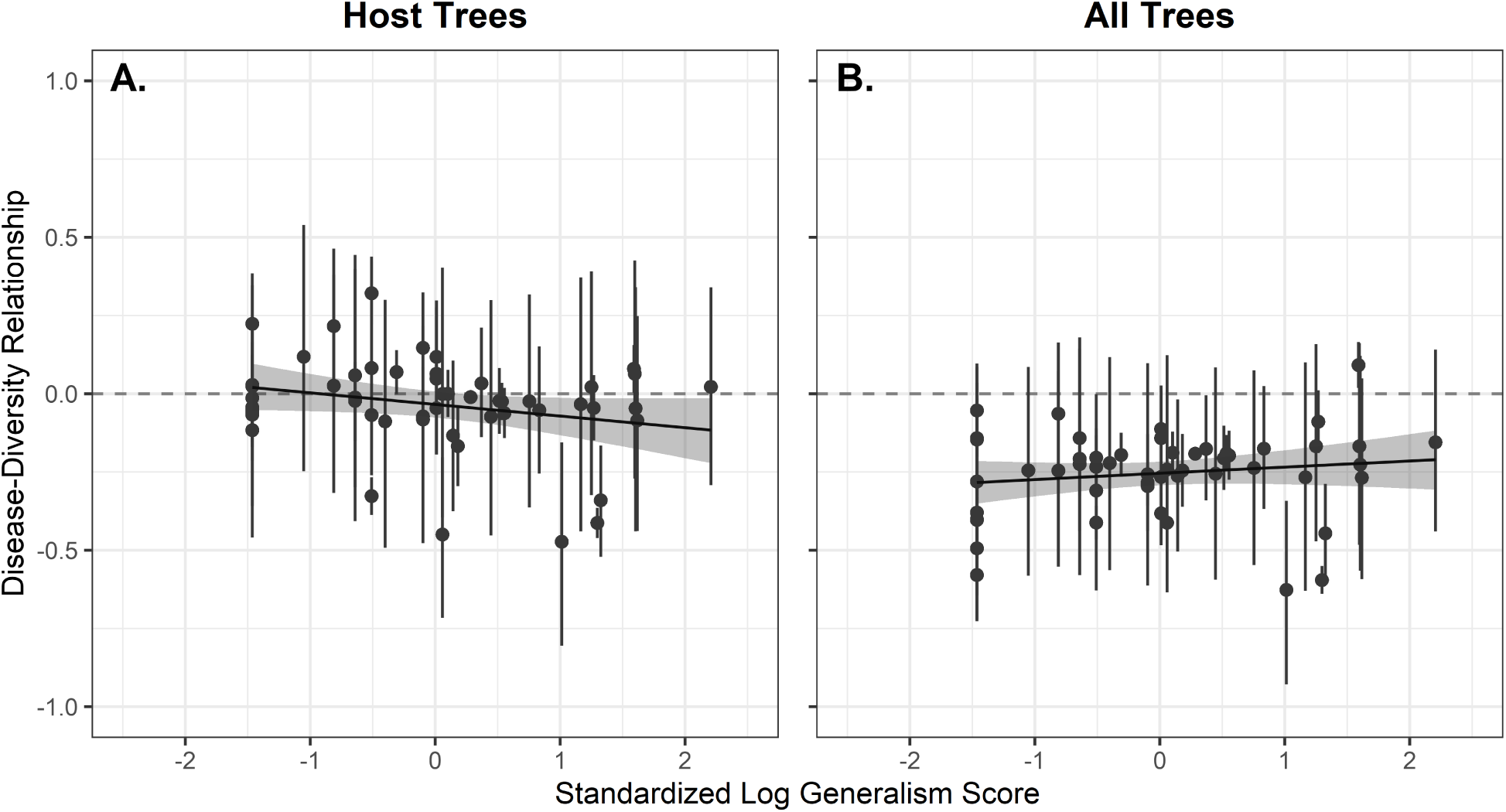
Relationship between pests’ disease-diversity relationships and their generalism scores when prevalence is assessed for either (A) host trees or (B) all trees. Pest disease-diversity relationships for each pest species are quantified as the posterior median slope between prevalence and community Shannon diversity. Error bars indicate 95% Bayesian credible intervals for each slope estimate. Black lines show fitted regression models and shaded regions show 95% Bayesian credible interval around the median trend.

To gain a preliminary understanding of whether specialist pests are subject to stronger dilution effects than generalist pests, we regressed the above-described species-specific disease-diversity relationships against two measures of pest species’ generalism. We quantified generalism for each pest using a global list of its known hosts at the genus level. To do so, we used either the total phylogenetic diversity of known host genera or a taxonomic measure which sums the number of each major taxonomic level present in the host list. As both measures give qualitatively similar results, we only report results for the taxonomic measure, which we call the generalism score. When we assessed prevalence for known hosts, we found a weakly negative relationship between generalism and the DDR indicating that specialists tend towards *weaker* dilution effects than generalists (Fig. 1A). Conversely, when we assessed prevalence across all tree species, this relationship was weakly positive, indicating specialists tend towards *stronger* dilution effects than generalists (Fig. 1B).

A simple regression as we have done above does not correctly propagate uncertainty in the estimated species-specific slopes to the estimated regression coefficient. Thus, we next combine both steps from the above analysis into a single statistical inference. For the combined analysis, we fit a second Beta-Binomial model where, instead of fitting species-specific slopes, we model pest prevalence as a function of either measure of pest generalism, Shannon diversity, and their interaction. We refer to this model as the “Interaction model.” Here, a positive interaction effect would imply that generalists experience weaker dilution effects, or tend towards amplification effects, whereas a negative interaction effect would imply that generalists experience stronger dilution effects. We focus on the results using the taxonomic generalism score, although the genus-level phylogenetic diversity gives similar results (Table 1). Indeed, we found a negative interaction when we calculated pest prevalence using only host trees and a positive interaction when we calculated prevalence using all trees (Fig. 2). This aligns with our finding from the species-specific slopes model; namely, generalists tend more towards dilution when we assess prevalence across known host trees, whereas specialists tend more towards dilution when we assess prevalence across all trees (Fig. 2, Table 1).

**Figure 2:**
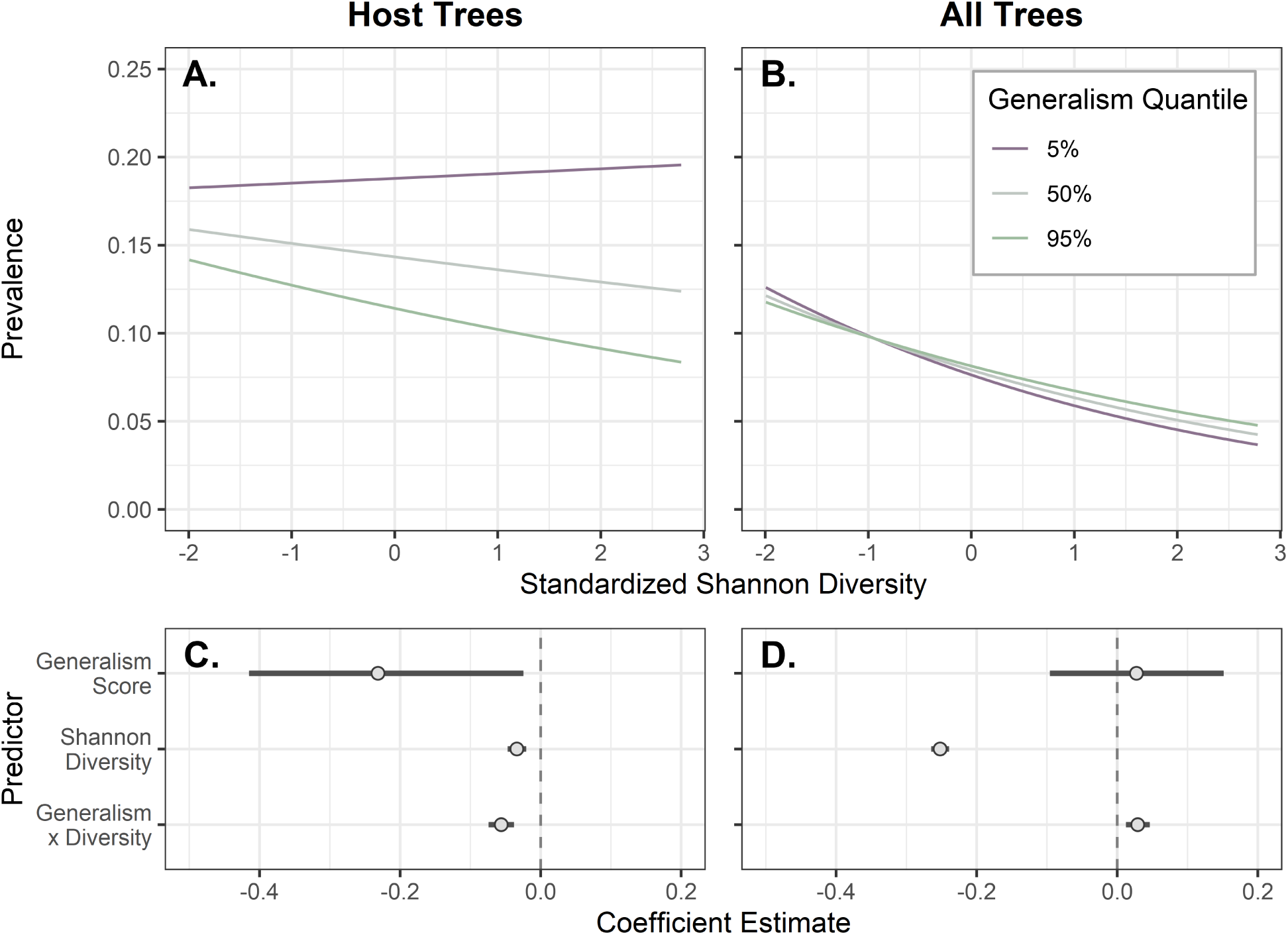
Interactive effects of plot Shannon diversity and pest generalism scores when prevalence is calculated for known host trees (left) or all trees (right). Panels A,B: Prevalence versus plot Shannon diversity (lines show 5%, 50%, and 95% quantiles for generalism values across all pest species in our data). Panels C-D: estimated model coefficients. Points indicate posterior medians and horizontal bars indicate 95% Bayesian credible intervals. Units of standardized measures represent one standard deviation in the Shannon diversity values across forest plots or generalism values across pest species. High generalism scores correspond to generalist pests while low scores correspond to specialists.

**Table 1:** Coefficient estimates and 95% credible intervals for the interaction model, using either the generalism score (Eq. 1) or genus-level phylogenetic diversity as a measure of pest generalism.

| Taxonomic Generalism Score |  |  |  |  |
|  | Host Trees |  | All Trees |  |
| Predictor | Median | 95% CI | Median | 95% CI |
|---|---|---|---|---|
| Generalism | -0.231 | (-0.415, -0.025) | 0.027 | (-0.096, 0.151) |
| Shannon Diversity | -0.034 | (-0.047, -0.021) | -0.252 | (-0.265, -0.239) |
| Generalism x Diversity | -0.056 | (-0.074, -0.038) | 0.029 | (0.012, 0.046) |
| Genus-level PD |  |  |  |  |
|  | Host Trees |  | All Trees |  |
| Predictor | Median | 95% CI | Median | 95% CI |
| Generalism | -0.237 | (-0.429, -0.040) | 0.010 | (-0.105, 0.130) |
| Shannon Diversity | -0.036 | (-0.049, -0.023) | -0.251 | (-0.264, -0.239) |
| Generalism x Diversity | -0.049 | (-0.067, -0.031) | 0.030 | (0.013, 0.046) |

### 2.2 Mathematical Model

To help understand our empirical results, we created a multi-host SIR epidemiological model. Specifically, we model a community containing *m* of *n* potential host species and a single pest species which is either a specialist or a generalist. For specialists, only one host is susceptible, whereas, for a generalists, all hosts are susceptible, albeit, with lower competence (Fig. 3 A-B). We ensure that the total susceptibility across the *n* host species (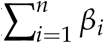, see Eq. 6 in Methods) remains the same for specialist and generalist pests. Thus, when we consider only host trees, a specialist pest encounters no variation in the susceptibility of its host species (because it only has one host species), however, it encounters high variation in the susceptibility of all tree species (compare bars labeled with *S* in Fig. 3A to those in Fig. 3 B). The generalist, on the other hand, encounters the same, moderate variation in host or tree susceptibility in both scenarios as all the trees are hosts (compare bars labeled with *G* in Fig. 3 A to those in Fig. 3 B). Because our empirical analyses showed that our estimated trends depended on how we calculated prevalence, we hypothesized that variation in susceptibility among the host community that we consider (i.e., host trees or all trees) may contribute importantly to differences in the strength of the dilution effect here as well.

**Figure 3:**
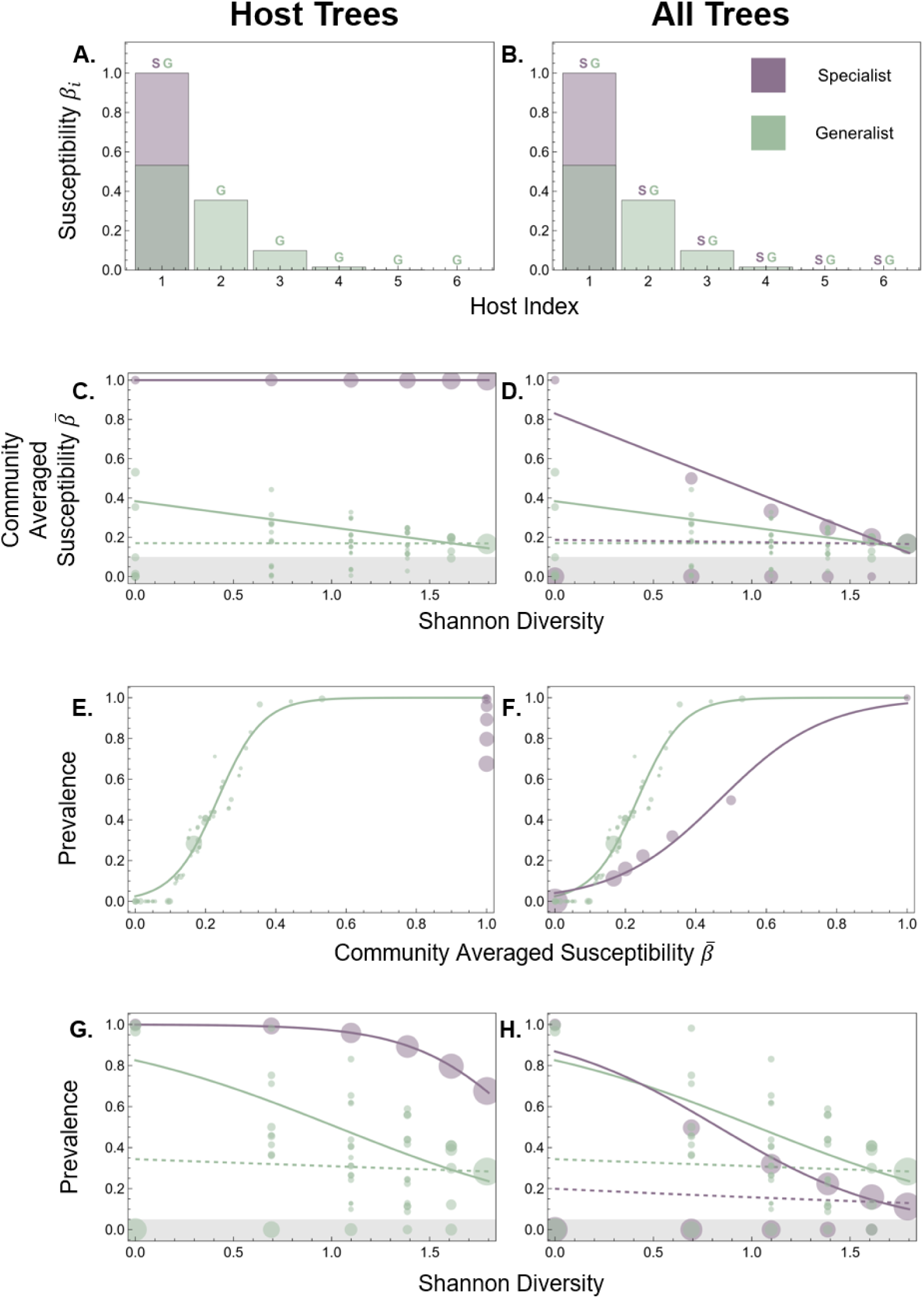
Disease-diversity relationships in the multi-host SIR model. For each level of species richness (*m ∈ {*1 to 6*}*), we generated 500 host communities at random. Circles represent communities in panels C-G, with the size representing the number of communities at a given point. Panels A-B: Susceptibility of tree species, *i ∈ {*1 to 6*}*, to the specialist and generalist pests. Note that for the specialist, susceptibility is zero for *i ∈ {*2 to 6*}*. Coloured letters above bars indicate whether the trees are included in community averaged susceptibility and prevalence calculations in the host trees (left column) and all trees (right column) cases for specialists (purple S) and generalists (green G). Panels C-D: Community averaged susceptibility when considering known host trees (C) or all trees (D) versus Shannon diversity of all trees in random communities. Lines show linear model fits either: excluding communities which fall below the *R*_0_ = 1 threshold necessary for spread, highlighted in grey, (solid); or including all communities (dashed). Note that the dashed and solid purple lines in C lie directly on top of one another. Panels E-F: Pest prevalence versus the community averaged susceptibility, when considering host trees (E) or all trees (F). Here, pest prevalence is calculated using the final epidemic size, and represents the proportion of the trees included in the calculation which are affected by a pest. Curves show fitted logistic models. Because all communities for the specialist in panel E have identical susceptibilities, we do not fit a purple curve in this case. Panels G-H: Pest prevalence, when considering host trees (G) or all trees (H) versus Shannon diversity of all trees in a community. Curves show logistic model fits either: excluding communities where the prevalence was zero, highlighted in grey, (solid); or including all communities (dashed). Note that the dashed and solid purple lines in G lie directly on top of one another.Parameters: *s*^Spec^ = 0, *s*^Gen^ = 0.1, and *γ* = 0.1. Model details are provided in the Methods section.

We use “final epidemic size,” defined as the total proportion of individuals infected over the entire course of the epidemic, to quantify disease prevalence. This quantity is dependent on the average susceptibility of the hosts in the community and, in our case, on both the pest specialization and the community composition. We produced variable community compositions for each possible species richness (*m ∈ {*1…*n}*) by randomly selecting *m* of the *n* potential host species from the regional pool, with replacement. For each community, we assumed that each host present was equally abundant.

The final size of an epidemic increases with increasing community-averaged susceptibility, *β* (Eq. 8), whether it is calculated as the frequency-weighted average susceptibility across only known host trees or across all trees in the community (Fig. 3 E-F). However, a dilution effect is observed even without a negative relationship between diversity and average susceptibility (Fig. 3 C-D and G-H). This is because final epidemic size depends on both average susceptibility and host abundance. Specifically, density dependent disease transmission results in a non-linear (concave) relationship between host abundance and final size. Species-poor communities may not include the host for a specialist pest, however, when they do, the host will be abundant (due to low host species richness). In contrast, species rich communities are more likely to contain a focal host, but at a lower abundance. Consequently, average realized susceptibility is constant across communities differing in Shannon diversity when we calculate susceptibility using only known hosts, but final epidemic size declines with species richness (Fig. 3 G, purple curve). For both specialists and generalists calculated either for known hosts or all hosts, we generate dilution effects of similar strengths (Fig. 3 G, H dashed lines; note the purple dashed line is identical to and, therefore, obscured by the purple solid line).

We do, however, find a difference in the dilution effect between the generalist and specialist pests in our model if we impose empirically-motivated nonrandom site selection. Specifically, studies of the DDR typically only consider communities where the pest or pathogen is present, as is the case in the dataset we have used here. In other words, biologists do not search for and make a record of pest prevalence in places where the final epidemic size is close to or even exactly equal to zero. To incorporate this into our model, we excluded communities which fall below the *R*_0_ = 1 threshold necessary for spread resulting in a pest prevalence of zero (i.e., we excluded the gray regions in Fig. 3). In doing so, we recreate the observed patterns in our empirical data (Fig. 2). Namely, we find that over intermediate ranges of community diversity, generalists exhibit a stronger dilution effect (more negative slope) than specialists when considering only known hosts (Fig. 3G) but a weaker dilution effect than specialists when considering all hosts (Fig. 3H). We note that this pattern holds when the community diversity is intermediate (Shannon diversity greater than *≈* 0.5 but less than *≈* 1.5, which is within the range present in our empirical dataset).

## 3 Discussion

Our analyses did not support the hypothesis that specialist pests are always more strongly diluted by biodiversity. Instead, our analyses reveal that that the strength and variation of an estimated dilution effect is sensitive to two often overlooked and important aspects of study methodology. Particularly, empirical and mathematical results regarding our hypothesis depended on whether prevalence was calculated for the entire community (all trees) or only across all the known hosts of a pest. Further, to reproduce the empirical finding in our epidemiological model, we had to impose a site selection bias by excluding sites with no evidence of a focal pest. Combined, this work both hints at a general mechanism for the DDR and demonstrates why observed dilution effects might differ qualitatively among studies, such that they should not be interpreted without considering the effect of these methodological considerations on the underlying data.

### 3.1 Variation in susceptibility may explain conflicting relationships between generalism and the DDR

Both the strength of an observed dilution effect and its relationship to pest host breath are sensitive to the measure of community-level prevalence. The overall strength of the dilution effect in the tree plot data was stronger when community-level prevalence was assessed for all species of tree than when it was when assessed only for trees known to act as hosts. We also observed a stronger dilution effect for specialists than for generalists when all trees were considered, but the opposite was true when only host species were considered. Importantly, we equate taxonomic measures of host breadth in our empirical test and variation in host susceptibility in our mathematical model to capture the key difference between specialists and generalists that could explain these results.

By comparing this result across both community-level prevalence measures and host breadths, the empirical results and mathematical models emphasize the importance of accounting for variation in the susceptibility of the species included in the prevalence metric when estimating a dilution effect. Considering that variation in host susceptibility is a necessary condition for the dilution effect to be observed (Keesing and Ostfeld, 2021), we hypothesize that more variation leads to stronger dilution effects, a pattern that is consistent with our results. In the case where all host species are included, a specialist encounters more variation in susceptibility among these hosts and, consequently, a stronger dilution effect compared to a generalist. Conversely, when only known host species are included, a specialist exhibits less variation among these hosts and, thus, a weaker dilution effect.

For known hosts, overall variation in susceptibility can still contribute to the dilution effect if transmission is density-dependent and if diluting non-hosts compete with hosts, thereby lowering their density. The strict specialist in our mathematical model still experienced a dilution effect when prevalence at the final size of the epidemic was calculated for its host alone. In our model set up, this is due to the decrease in density of host species in the presence of non-host species, coupled with density-dependent transmission. Similarly, holding density constant in cases of more generalist species allows us to directly interpret the community-averaged basic reproductive number as community competence. This interpretation also highlights the scope of our hypothesis. While the communities used in our mathematical model assume a constant total density as biodiversity increases, forest density may also increase with biodiversity and lead to amplification if some host interactions are mutualistic rather than competitive (Su et al., 2022). Additionally, the transmission of pests may be frequency-dependent rather than density-dependent, leading to different dilution effect mechanisms (Toorians et al., 2025). Therefore, deviations from our model assumptions in real ecosystems could help explain why our empirical result, but not our mathematical model, suggests that we tend to observe amplification effects for specialists in the hosts-only case.

### 3.2 Site-selection bias may confound dilution effects attributed to naturally-occurring relationships between biodiversity and community competence

In addition to life-history theory explanations and real biological relationships between biodiversity and community competence (Huang et al., 2013; Joseph et al., 2013; Halliday et al., 2020), site selection biases can also contribute to the biodiversity-competence relationship and, hence, the strength of the dilution effect. Imposing non-random site selection in our mathematical model both introduced a negative biodiversity-competence relationship and led to a stronger dilution effect. This supports idea that a negative biodiversity-competence relationship is an important contributor to the dilution effect. This general observation is consistent with previous studies comparing dilution effects for random biodiversity gradients with non-random biodiversity gradients which produce negative biodiversity-competence relationships (Halliday et al., 2020; Johnson et al., 2013; Rosenthal et al., 2021). That being said, our mathematical model reveals that this negative biodiversity-competence relationship can also arise simply due to study design. Furthermore, in the case when all trees were considered in the measure of prevalence, the dilution effect increased in strength more so for specialists than for generalists, though there was no change for specialists in the host trees case. This suggests that site selection bias may impact estimated trends for specialists and generalists differently and that this also depends on the measure of prevalence considered.

### 3.3 Limitations and future directions

While we focus on host regulation and the relationship between biodiversity and community-averaged competence as the main mechanisms driving the dilution effect, other mechanisms are also expected to contribute. For example, encounter reduction and the resource concentration hypothesis (Keesing and Ostfeld, 2021; O’Rourke and Petersen, 2017) depend on how the interactions between hosts and pests change in the presence of other associated species, separate from the direct effects of density or frequency. For insect pests, the presence of some plants might hinder dispersal to suitable hosts, decreasing transmission. These mechanisms, however, are likely to depend on specific pest and host species and would be better studied at the species, rather than community, level. While these and other alternative mechanisms may contribute to the dilution effects found in our empirical data our analysis cannot test for them specifically (but see Rosenthal et al., 2021) and they are not present in our mathematical model. To our knowledge, there is limited use of multi-host epidemiological models to investigate hypothesized dilution effect mechanisms. Future work formalizing verbal dilution effect theory into multi-host models under different modes of transmission and ecological assumptions will be important (Toorians et al., 2025).

In designing our mathematical model, we also made several simplifying assumptions about the general epidemiological, demographic, and eco-evolutionary mechanisms that could influence the dilution effect. The mathematical model assumes density-dependent, not frequency-dependent transmission. By equating empirical results from recorded damage to the final epidemic size of an SIR model, we assume that damage from pests is permanent and that outbreaks are complete. Alternatively, an SIS model could reflect equilibrium conditions if instead we assumed that infected hosts return to a susceptible state on a comparable time scale to infection dynamics. Generally, the most appropriate epidemiological model will depend on the exact pest species. The epidemiological model also omits host demography and pest virulence in order to set specific community compositions and transmission rates. In real ecosystems, we would expect demography to change the community composition through time and to depend on which pests and their hosts are present in the community. Additionally, our randomly assembled communities do not necessarily reflect realistic community compositions.

While we focus on community-level prevalence, there are many ways to quantify disease or pest impact when considering their relationships to biodiversity. Our measure of pest prevalence that includes all trees is a host-centric measure and may be applicable in cases when all or most of the tree species in a community are of interest, such as forest management. Including only the host trees in prevalence is a more pest-centric metric which, instead, measures the success of a pest within its niche. In more targeted applications, considering prevalence per species of host is important when we are interested in the impact of a pest or pathogen on a selected species of host. For example, work on zoonotic or vector-borne diseases such as Lyme disease may focus on infection prevalence in human hosts or in particular disease vectors (e.g. Ostfeld and LoGiudice, 2003). Rather than focusing on one pest at a time, prevalence can also be aggregated across all pests to assess overall pest or disease burden. Infestation, disease, or damage severity could all be assessed at the level of host individuals as well (e.g. Jactel et al., 2021). Although studies which use these all fall under the broad category of disease-diversity relationships, it is important to distinguish them to isolate underlying mechanisms and applied interpretations (Rosenthal et al., 2021; Roberts and Heesterbeek, 2018).

Our study highlights the importance of two key methodological considerations when studying the dilution effect. First, the data and calculations used to quantify pest impact or prevalence is crucial. We find differences in results even between similar community-level prevalence measures and expect to see differences across the many other ways pests and disease can be measured. We suggest dilution effect studies should test and report multiple measures with the same data where relevant and, at minimum, clearly describe the chosen measure to facilitate comparison to other studies. Second, site-selection bias, which is likely present in many field studies, can introduce a confound that potentially increases the strength of an observed dilution effect. By considering these factors, further analyses and models may resolve contradictory results in the literature to help answer important applied questions surrounding the generality and management of the dilution effect.

## 4 Methods

### 4.1 Data and Calculations

We used pest incidence data from Gougherty and Davies (2024), which was derived from the US Forest Service Forest Inventory Analysis (**?**U.S. Department of Agriculture, Forest Service, 2025). Pest taxa comprised primarily insects and fungi, with one parasitic plant. Pests were not detected directly but, instead, inferred to be present when there was visible host damage known to be caused by that pest. For each pest species detected in a plot, the derived dataset includes the number of trees with recorded damage from that pest and the total number trees that are known hosts of that pest. Plot-level data in the derived dataset includes the Shannon diversity and total number of trees in a plot. After filtering these data to include only pests which were present in at least two plots, our dataset comprised 52 pest species found across 26549 plots. As multiple pests could be present in the same plot, this resulted in a total of 30719 pest-plot pairs.

To assess generalism for each pest species, we obtained the global list of known pest-host associations used in the study by Gougherty and Davies (2024). While the majority of these records are at the species level, we chose to assess generalism at the genus level in order to help account for differences in pest species’ ranges; pests with large range sizes may associate with multiple species of hosts in different parts of their range that are ecologically similar. Alternatively, some pest species might be partially delineated by host, artificially creating more specialist species. Assessing generalism at the genus level also avoids issues with taxonomic changes and poorly resolved or incomplete phylogenetic trees available at the species level.

Generalism for a pest or pathogen can be treated categorically (specialist versus generalist) or quantitatively by using some biodiversity measure of its hosts (Poulin et al., 2011). Our main generalism metric is a modified version of a ”taxonomy as phylogeny” generalism score from Lewthwaite et al. (2018) (Eq. 1).

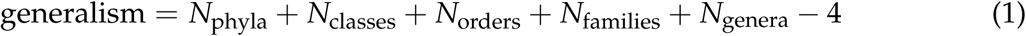

This taxonomic diversity quantifies the idea that pests that attack hosts from more, and more distant taxa are more generalist, while accounting for the fact that having a host at one of any taxonomic level implies there is least one of each of the other taxonomic levels. This measure makes no claim about the rate of host switching, and uses taxonomy as a measure of relative phenotypic differences among hosts, ignoring intrageneric differences (see above). Our secondary metric makes use of a dated phylogeny linking hosts and so is an explicit model of gradual phenotypic change. We first amalgamated a dated genus-level phylogenetic tree of hosts from multiple sources and then calculated generalism as the total phylogenetic diversity (PD, sensu Faith, 1992) of hosts, i.e. the sum of the branchlengths (in millions of years) connecting the genera used by a pest. The majority of the phylogeny was from a recent genus-level study of angiosperms and gymnosperms (Dimitrov et al., 2023). We added fern genera using a fern phylogeny from Nitta et al. (2022), and used the divergence time of the two genera of lycophytes in Testo et al. (2018), supplementary figure 4. We used the divergence times between lycophytes, ferns, and spermatophytes as estimated in Harris et al. (2022). We note that there is a great deal of variation in patristic distances among hosts (and their associated errors) on this deep phylogeny, and thus large variation in this PD-based measure of breadth, so we prefer our simpler, taxonomic measure.

For both measures of generalism, we used the rgbif package in R to obtain higher taxonomic classifications for each unique host species of genus in the list (Chamberlain et al., 2026). We resolved synonyms, ambiguous species names or taxonomy that did not match the phylogenetic tree using records in The Integrated Taxonomic Information System (ITIS) or The Catalogue of Life (Integrated Taxonomic Information System, 2026; Bánki et al., 2026).

### 4.2 Statistical Analysis

Using a modified version of the community-level disease-risk model presented in Rosenthal et al. (2021), we model the number of hosts affected by species *s* in plot *p*, which we denote as *Y_s_*_,*p*_, using a Beta-Binomial regression where the number of trials is either 1) the number of known host trees for the pest in the plot or 2) the total number of trees in the plot. Note that, following Gougherty and Davies (2024), we only model species *×* plot combinations for which there is at least one detected parasite of species *s*. Specifically, we model *Y_s_*_,*p*_, as:

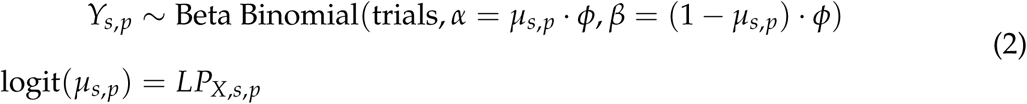

Here, *ϕ* denotes a dispersion parameter that measures the amount of dispersion compared to a standard binomial model (*ϕ* = 0 corresponds to a standard binomial model), and *LP_X_*_,*s*,*p*_ denotes a linear predictor model for the mean probability of trees being affected, *µ_s_*_,*p*_. We use *X* as a place-holder here, as we consider two different models with this term, described next.

For each of the two prevalence scenario described above, we consider two different linear predictor models. First, in what we call our species-specific slopes model (denoted by SS), we estimate the effect of the Shannon diversity of trees in plot, *p*, denoted by *H_p_*, on prevalence for each pest species, *s*, fitting a separate intercept, *a_s_*, and slope, *b_s_*, for each. Specifically,

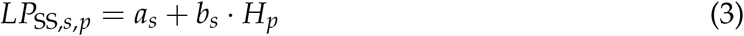

We subsequently perform a linear regression to quantify the relationship between a pest species’ slope, *b_s_* and its generalism score, *G_s_* to assess the strength of a potential disease-diversity relationship. We fit the model

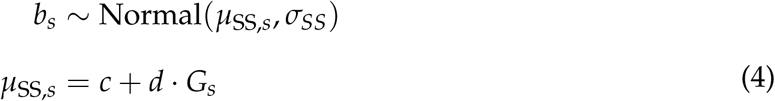

where *d* quantifies the relationship (slope) between the DDR and generalism, and *µ*_SS,*s*_ is the DDR for pest species *s* estimated using species-specific slopes model.

The above approach does not account for differences in the species’ sample sizes (each species was detected at a potentially different number of plots) and, therefore, error in both estimated coefficients *c* and *d* does not correctly reflect uncertainty in the estimated species-specific slopes. Thus, we run a second model, which we call our interaction model (denoted by INT). Here, we model the interactive effects of two independent variables, Shannon diversity of plot *p*, *H_p_*, and generalism of species *s*, *G_s_*. Specifically,

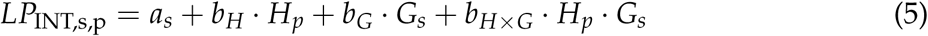

Because generalism is measured at the pest-species level, we do not include species-specific slopes in this model. Instead, we let the interaction term fill a similar role, as the interaction modifies a species’ slope based on that species’ generalism score. While this second model is not as flexible (i.e., species’ slopes are only able to differ based on the deterministic effect of differences in their score), a model that includes an interaction as above as well as species-specific slopes will have any differences in species’ slopes absorbed into the each species’ specific slope term, leaving little variation to be explained by the interaction term.

We fit all statistical models in a Bayesian framework using Stan (Stan Development Team, 2025). Measures of generalism were log-transformed, and both generalism and Shan-non diversity were standardized to a mean of 0 and standard deviation of 1 before model fitting. We used r-hat*<*1.01 and bulk-ESS*>*400 to assess model convergence for 4 chains. To achieve this, we run the linear regression and interaction model for 3000 iterations, and the species-specific slopes model for 6000 iterations.

### 4.3 Mathematical Model

We created a multi-host epidemiological model to better understand our empirical results. The model included *n* = 6 host species in a regional pool which we allowed to differ in their susceptibility to a pest, as variation in host competence (susceptibility or onward transmission) is a necessary condition for the dilution effect. We do so by varying the transmission rate, *β_i_*, for *i* = 1, . . . , *n* in the model for each host. We explicitly define a specialist pest and a generalist pest in such a way that host breadth is connected to the variation in susceptibility. Here the specialist can utilize only a single host, while the generalist can utilize any host (*i*) to differing degrees. To generate pests with differing host breadths we utilize a binomial distribution to define the pest’s set of host-specific transmission rates, *β_i_* (Fig. 3 A, B:

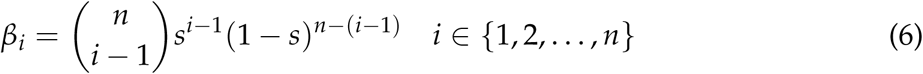

The parameter *s* determines the host breadth of a given pest: the strict specialist is able to infect a single host species as given by 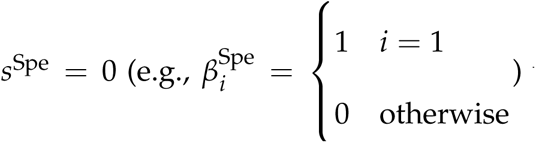 whereas the generalist has a non-zero value of *s*^Gen^ *>* 0 (hence 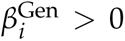 for all *i*). This method of defining host breadth assumes that the total host susceptibility 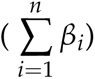 is equal for all pests, but is distributed among potential host species differently based on *s*. When *s* is close to zero, the pest is most specialized on one species of host.

As with our empirical analysis, we considered pest prevalence across all possible hosts in the community, or only the known hosts of the pest of interest. Let *H* be the set of all hosts in the community and *H̊* the set of all known hosts of a focal pest. Here we choose the set of known hosts to reflect all hosts with a strictly non-zero susceptibility to the pest (*H̊* Spe = *{*1*}* and *H̊* Gen = *H* = *{*1, 2, . . . , 6*}*). We hypothesize that the realized variation in susceptibility among hosts of a focal pest, whether that variation in measured among all hosts *H* or only among known hosts *H̊* , can explain the strength of the disease diversity relationship. In our scenarios, the specialist shows no variation in susceptibility when considering only its recognized host, but high variation in susceptibility when considering all potential hosts. The generalist, on the other hand, shows an intermediate level of susceptibility regardless of how prevalence is measured. Hosts are assumed to differ only in transmission rates in regard to susceptibility to the focal pest, not onward transmission nor their recovery rates (*γ*).

Defining the susceptibilities of individual hosts to a focal pest, we can model the resulting dynamics of a focal parasite within the multi-species community using a classic susceptible (S), infected (I), recovered (R) compartmental model:

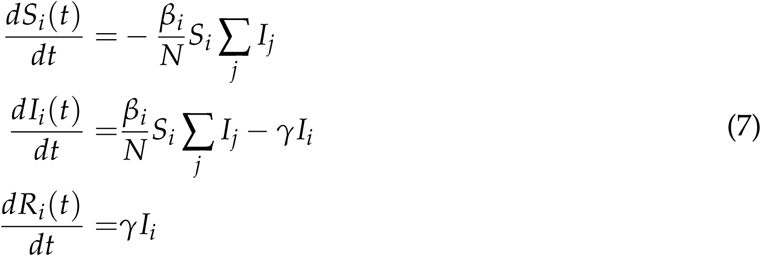

There are several possible approaches to generating ”random” host communities with varying frequencies of hosts of type *i*, denoted *h_i_*, given a set total number of individuals. Here, we generate random communities by first specifying species richness *m ≤ n* of a community, randomly selecting *m* of the *n* possible hosts then assuming each host is equally common (i.e., *h_i_* = <u>^1^</u> if host is present and *h_i_* = 0 otherwise). We quantify the resulting diversity of the community using the Shannon index (∑*_j∈H_ N_j_* ln(*N_j_*) where *N_i_* = *h_i_ N*). Note that Shannon diversity is always measured among all hosts in the community. In this model, the total size of the host community remains constant and as given by *N* = ∑*_j_*(*S_j_* + *I_j_* + *R_j_*), and the frequencies of each host also remain constant as defined using the approach above.

Epidemiological dynamics depend on both the host susceptibilities and their relative frequencies, and can be summarized as a community averaged measure of susceptibility and the community basic reproductive number. Let *h̊_i_* denote the frequency of host *i* among the subset of hosts considered (e.g., among known hosts 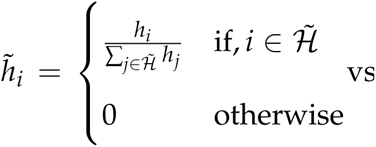vs. among all hosts *h̊_i_* = *h_i_∀i*). We can then define the ”community averaged susceptibility” (*β*) of a community as:

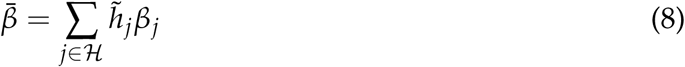

When considering only known hosts, the average susceptibility of hosts to the specialist parasite is independent of host community diversity (Fig. 3 C), as the realized frequency of the focal host (i.e., host 1) is constant, *h̊*_1_ = 1. In contrast, for generalists or specialists when considering all possible hosts, the average susceptibility of communities varies but is on average independent of Shannon diversity (Fig. 3D). In this SIR model, a pest will spread if the community basic reproductive number is greater than 1.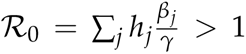 This is the average of the individual basic reproductive numbers for each host, *R*_0,*i*_ = *β_i_*/*γ* weighted by the frequency of each host type in the community. Given the pest spreads, this SIR model exhibits classic epidemic dynamics with the disease first increasing in frequency before ultimately reaching heard-immunity and then dying out.

As an analogue to pest prevalence in the empirical data, we follow (Springbett et al., 2003) and quantify disease severity in terms of the ”final epidemic size”, denoted *Y_∗_*_,∞_, which quantifies the proportion of all hosts infected at the end of the epidemic. The final size of an epidemic in a multi-host community is given by the transcendental equation:

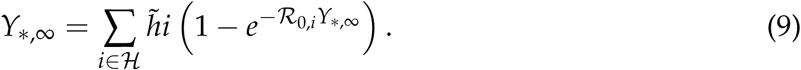

where the (*∗*) indicates that this proportion can be measured among all hosts or only among known hosts. We then compare these to Shannon diversity of the community to assess the disease-diversity relationship.

## 5 Acknowledgments

The authors thank A. V. Gougherty and Z. Wang for providing supplementary datasets and information related to the data used and T. J. Davies for insightful discussion during the conceptualization of this work. We also thank members of our research labs for their comments and feedback.

